# Layered structure and complex mechanochemistry of a strong bacterial adhesive

**DOI:** 10.1101/183749

**Authors:** Mercedes Hernando-Pérez, Sima Setayeshgar, Yifeng Hou, Roger Temam, Yves V Brun, Bogdan Dragnea, Cécile Berne

## Abstract

While designing adhesives that perform in aqueous environments has proven challenging for synthetic adhesives, microorganisms commonly produce bioadhesives that efficiently attach to a variety of substrates, including wet surfaces that remain a challenge for industrial adhesives. The aquatic bacterium *Caulobacter crescentus* uses a discrete polar polysaccharide complex, the holdfast, to strongly attach to surfaces and resist flow. The holdfast is extremely versatile and has an impressive adhesive strength. Here, we use atomic force microscopy (AFM) to unravel the complex structure of the holdfast and characterize its chemical constituents and their role in adhesion. We used purified holdfasts to dissect the intrinsic properties of this component as a biomaterial, without the effect of the bacterial cell body. Our data support a model where the holdfast is a heterogeneous material composed of two layers: a stiff nanoscopic core, covered by a sparse, flexible brush layer. These two layers contain not only *N*-acetyl-*D*-glucosamine (NAG), the only yet identified component present in the holdfast, but also peptides and DNA, which provide structure and adhesive character. Biochemical experiments suggest that, while polypeptides are the most important components for adhesive force, the presence of DNA mainly impacts the brush layer and initial adhesion, and NAG plays a primarily structural role within the core. Moreover, our results suggest that holdfast matures structurally, becoming more homogeneous over time. The unanticipated complexity of both the structure and composition of the holdfast likely underlies its distinctive strength as a wet adhesive and could inform the development of a versatile new family of adhesives.

---

Industrially produced adhesives pale in comparison to the versatility and compliance exhibited by the diverse biological adhesives available from nature. Many organisms, from microscopic bacteria to larger vertebrates, are able to attach to surfaces, using strategies based of the production of elaborate adhesive biomolecules or complex physical structures. For example, mussels and barnacles produce a multi-protein complex that acts as a wet adhesive to attach to various surfaces ^1,2^. On the other hand, geckos toe pads are composed of a hierarchical structure of lamella consisting of thousands of micron-sized setae. Each seta is made of hundreds of nano-scale spatulas that mediate strong attachment via van der Waals and capillary interactions ^3^. But the widest diversity of bioadhesives is produced by microorganisms.

Due to their small size, bacteria have evolved nanoscopic bioadhesives to colonize most surfaces. Some of the best-studied bacterial adhesins are proteinaceous and include long polymeric structures, such as fimbriae and pili, and short non-fimbrial adhesins, all of which are directly anchored to the cell surface ^4^. Recent biophysical studies of bacterial protein adhesins, conducted by Atomic Force Microscopy (AFM), have led to models whereby the concerted action of multiple relatively weak adhesion molecules ^56^, or micro-domain accumulation of adhesins at the cell-substrate interface ^7^, can mediate strong bacterium-substrate interactions. Another emerging theme is that multi-domain protein adhesins can mediate versatile binding to different substrates ^8-10^. Polysaccharides are another important class of bacterial adhesives. While there is a large body of work on the mechanochemical properties at the single molecule level of single polymeric polysaccharides ^11,12^, only few studies investigate the physicochemical properties of bacterial polysaccharide adhesives ^9,13,14^

One of the best-studied bacterial polysaccharide adhesive is the unipolar polysaccharide adhesive (UPP) produced by multiple genera of the alphaproteobacteria ^4,15,16^. The prototype UPP adhesive is the holdfast produced by the oligotrophic fresh water bacterium *Caulobacter crescentus* (Fig. 1 A, B), commonly found in aquatic environments low in nutrients such as pristine lakes, tap water supplies or distilled water tanks ^17-19^ The holdfast adhesive is extremely versatile, being able to adhere to a variety of biotic and abiotic surfaces ^4,20^, and exceptionally strong, with a tensile strength exceeding 68 N/mm^2^, one of the highest measured for both synthetic and biological adhesives ^21^. These properties make the holdfast a promising bioadhesive for a variety of applications ^22^. The holdfast is a small, elastic ^23^ structure located at the end of a long, thin extension of the cell envelope called the stalk (Fig. 1 A, B). In natural environments, the stalk places the cell body well within the flow and away from the stagnant substrate boundary layer, allowing improved access to nutrients in typically oligotrophic conditions ^24^. The tradeoff for this adhesion strategy is that large drag forces on the cell body have to be balanced by an anchor with a nanoscopic footprint (Fig. 1C). The use of a nanoscopic footprint rather than an adhesive distributed over the whole cell surface suggests that the holdfast might be a material of composition and structure distinctly different from those of previously studied bioadhesives.

**Fig. 1:**
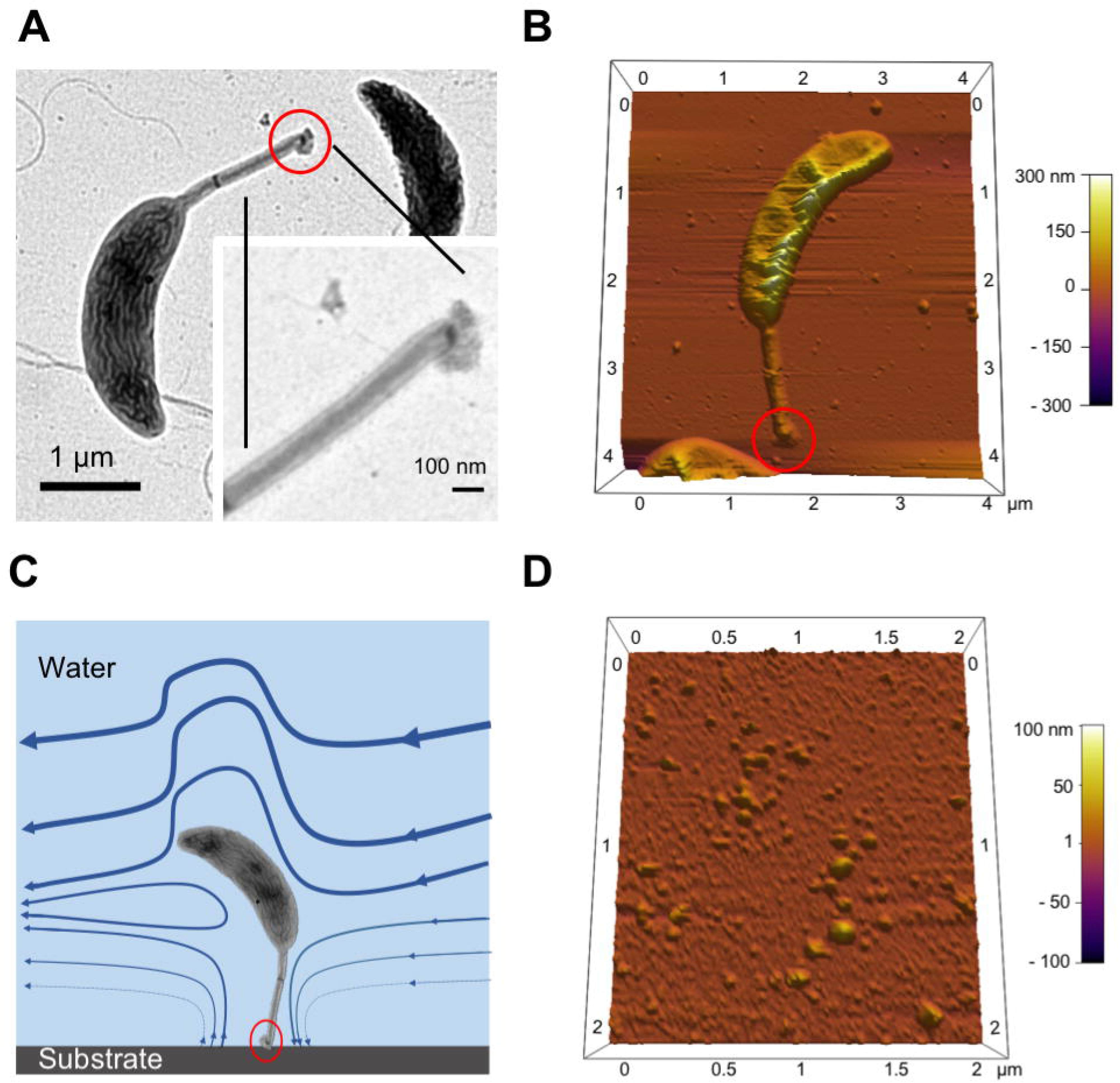
Caulobacter crescentus. (A-B) TEM (A) and AFM (B) images of a *C. crescentus* CB15 wild-type cell. (C) Schematic of *Caulobacter* adhesion on surface via the holdfast located at the tip of the stalk under fluid flow. The holdfast size is in the order of tens of nanometer in diameter. Holdfast is circled in red. (D) AFM image of purified holdfasts in dH_2_O.

Determination of holdfast morphology and composition is difficult, due to its high adhesiveness, insolubility, and the small amount (less than 10^−3^ μm^3^) produced by each bacterium. The only known component of the holdfast is a polymer of *N*-acetyl-*D*-glucosamine (NAG) ^25^, but its composition is probably more complex, to achieve such a strong adhesion. Electron microscopy studies ^19,20,25,26^ and AFM imaging ^23,27,28^ suggest that the holdfast is amorphous. Indeed, newly secreted holdfast appears to spread over the surface as a viscous fluid that subsequently hardens ^27^. Consistently, single molecule force spectroscopy (SMFS) studies conducted on purified holdfasts showed that holdfast adhesion is time-dependent, and adhesion force strengthens quickly over time ^28^. This study also suggested the presence of discrete, diffusible adhesins of unknown nature, responsible for the bulk of adhesion ^28^.

Here, by studying purified holdfasts to avoid any mechanochemical contribution of the stalk and/or the cell body (Fig. 1D), we show that holdfast adhesive properties stem from a complex structural organization and composite mechanochemistry. Using SMFS, we find that the holdfast is composed of two different layers with distinct physicochemical properties: a stiff, solid-elastic core, covered by a sparse biopolymeric brush layer. Enzymatic treatment experiments show that peptides and DNA molecules are important for the integrity of the brush layer, while NAG polymers seem to be mostly present in the core. Finally, dynamic adhesion force spectroscopy experiments show that peptides are crucial for holdfast strength of adhesion, while NAG residues predominantly play a structural role, and DNA is involved in initiating holdfast adhesion. We describe the holdfast as a bioadhesive combining a complex multilayer organization and diverse chemistry. Bioinspired adhesives have already been developed based on the mussel DOPA wet adhesives and physical structures of the gecko seta ^29-31^, suggesting that further study of complex adhesives such as the holdfast can inform the development of even more versatile adhesives.

## Results and Discussion

### Mechanical Characterization of the Holdfast by AFM Indentation Experiments

#### Holdfast core elasticity

The holdfast initially displays fluid-like properties, wetting the substrate upon contact within minutes ^27^. However, strong adhesion requires a viscoelastic or elastic-solid adhesive. To determine whether the holdfast acquires elastic-solid properties over time, we measure the deformation of holdfast over time under a constant loading force and perform creep compliance experiments ^32^ on holdfasts deposited on a freshly cleaved mica surface 16 h before measurements. In these experiments, the deformation changes are measured *vs*. time under constant load ^33^. After applying a constant loading force of 40 nN, holdfasts do not exhibit measurable creep over a dwell time of 120 sec (Fig. 2A). If strain were observed in time under constant stress, the material would have qualified as plastic or viscoelastic. Instead, at least over an experimental time scale of minutes, cured holdfasts respond mechanically as a solid elastic material.

**Fig. 2.**
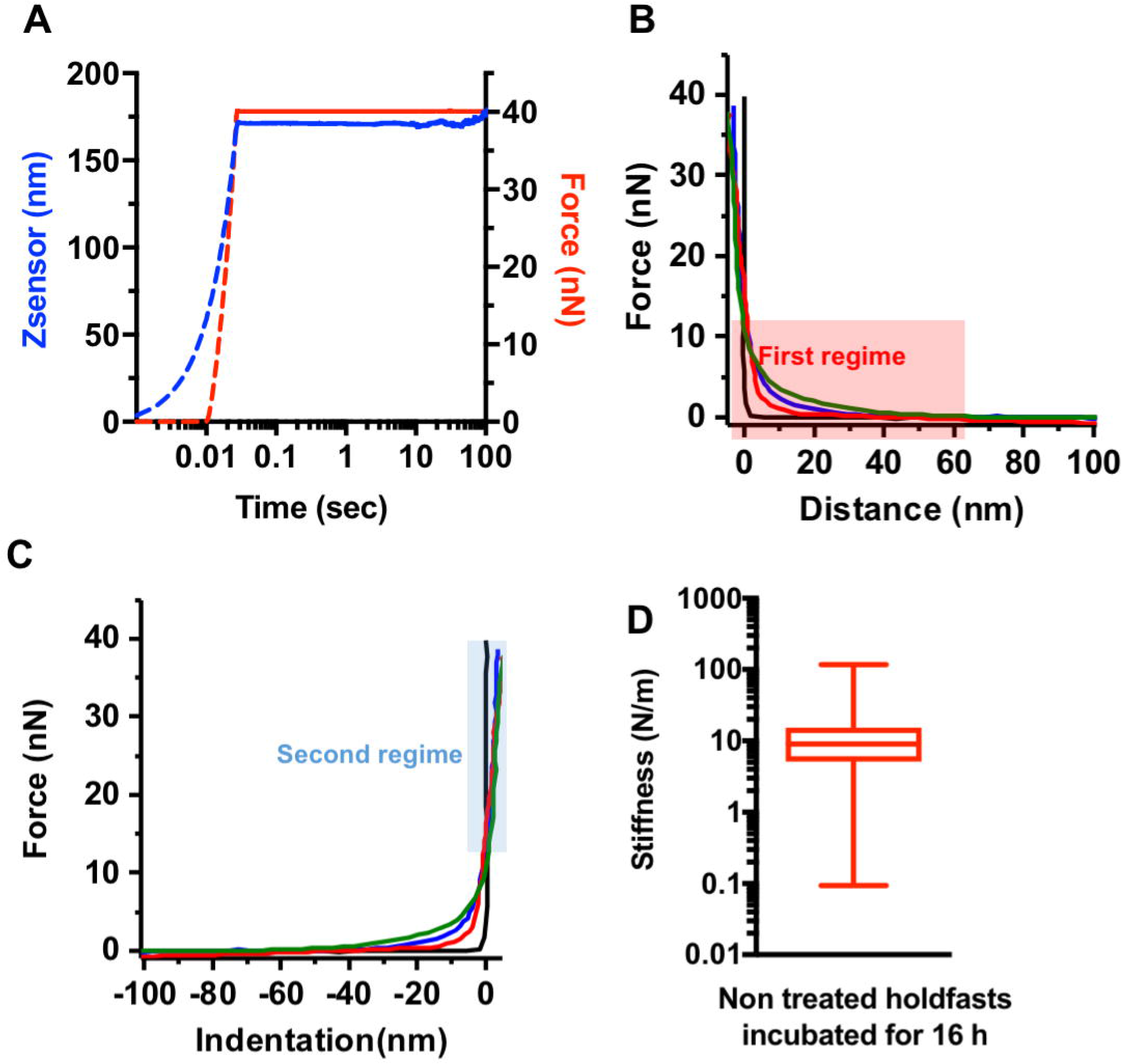
Holdfast core and biopolymer brush layer characterization. Representative creep compliance experiment performed on a holdfast with a constant force of 40 nN with a dwell time of 120 sec. First, the AFM tip indents the holdfast particle until the trigger force is reached (dotted curves). Once the probe reaches the set trigger force (40 nN), the particle is kept at a constant force for 120 sec (solid curves). Particle deformation over time is recorded as changes in piezo extension Z (Zsensor channel, blue curve) and translated to Force (red curve). (B-C) Examples of force *vs*. distance (B) and force *vs*. indentation (C) curves performed on different holdfasts (blue, red and green curves) or the mica substrate (black curve). The first (far from the surface) and second (close to the surface) regimes are depicted as red and blue boxes respectively. (D) Box and whisker plot of stiffness of 16 hours cured holdfasts, calculated from the slopes of the linear part of the second regime of curves shown in panel (B).

Next, we perform AFM compressive stress/strain (indentation) experiments. Fig. 2B and 2C show force *vs*. distance (F *vs*. D) and equivalent force *vs*. indentation (F *vs*. Ind) curves respectively on three different holdfasts and on clean mica as a control. Those curves reveal two clearly different interaction regimes between AFM tip and holdfast. The first regime can be characterized as a soft, repulsive interaction, manifested over a remarkably long range (> 60 nm) (Fig. 2B, red box). The second regime is characterized by a much stiffer response (Fig. 2C, blue box).

We first focus on the second regime, given by steep slopes in the F *vs*. Ind curves, corresponding to the innermost layer or core of the holdfast. Since compression creep compliance tests reveal a behavior consistent with a solid, elastic body (Fig. 2A), an effective spring constant of the holdfast core is obtained from the slopes of the approximately linear part of indentation curves (Fig. 2C and Fig. S1). The standard deviation (SD) from linear fit slopes is less than 2% for each analyzed curve. We observe a broad distribution of apparent stiffness of 9.0 ± 12.6 N/m (median ± SD) (Fig. 2D). In addition, the wide distribution of data suggests a significant heterogeneity, both at the single holdfast level and between holdfasts (Fig. S2A). The measured stiffness is extremely high for a bioadhesive and is 1-2 orders of magnitude higher than that measured on exopolysaccharides (EPS) or lipopolysaccharides (LPS) present on the surface of bacteria ^34^.

Since the tip/sample area increases in principle while indenting, the dependence of force on indentation is nonlinear and described by the Hertz model for small indentations (Supplementary information). Application of this model to the stiff response portion of F *vs*. Ind curves, corresponding to the second regime, allow estimating the average Young’s modulus, E, of the core material of 0.37 ± 0.18 GPa (Table 1), comparable to that of amyloid fibers ^35^, which are involved in adhesion of different bacteria ^36^ or of mussels byssal threads ^37^), but one and two orders of magnitude higher than the EPS from *Staphylococcus epidermidis* ^14^ and *Pseudomonas aeruginosa* ^38^, respectively.

Previous work ^39^ has analyzed fluctuations in stalk angle for a pair of whole cells attached to a single holdfast, measuring an effective torsional spring constant from which an elastic modulus, *E* ≈ 2.5 × 10^4^ Pa, is determined. This value is approximately 3 orders of magnitude smaller than that obtained by direct compression measurements of purified holdfast reported here. A possible explanation for this discrepancy is that the torsional spring constant obtained from whole cell measurements likely reflects the shear modulus of the anchor proteins that connect the holdfast to the stalk in addition that of the holdfast itself. We note that the simple form of the Hertz model assumes that the sample is homogeneous, isotropic and infinitely thick, whereas real biological samples deviate from these idealized conditions. Recent studies have investigated the validity of these assumptions by explicitly testing the dependence of the elastic modulus on indentation depth ^40^. Fig. 3A-C shows numerical simulations of the strain field for a two-dimensional layer of elastic material with Young’s modulus representative of holdfast samples on a noncompliant substrate, indented by an AFM tip of radius 13 nm. We note that the strain field extends several times the tip radius into the sample. While the deformation is consequently thickness-dependent, the force-indentation response curves converge beyond this threshold of thickness for small indentations (Fig. 3D). Given measured holdfast thicknesses in the 20-100 nm range by AFM imaging (Fig. S3), finite thickness corrections to the simple Hertz model are also included ^41,42^ (Supplementary information). The results of fits with and without finite thickness corrections are consistent (to within a factor of < 2), demonstrating that under our experimental conditions the substrate does not impact the order of magnitude of the measured holdfast stiffness.

**Fig. 3:**
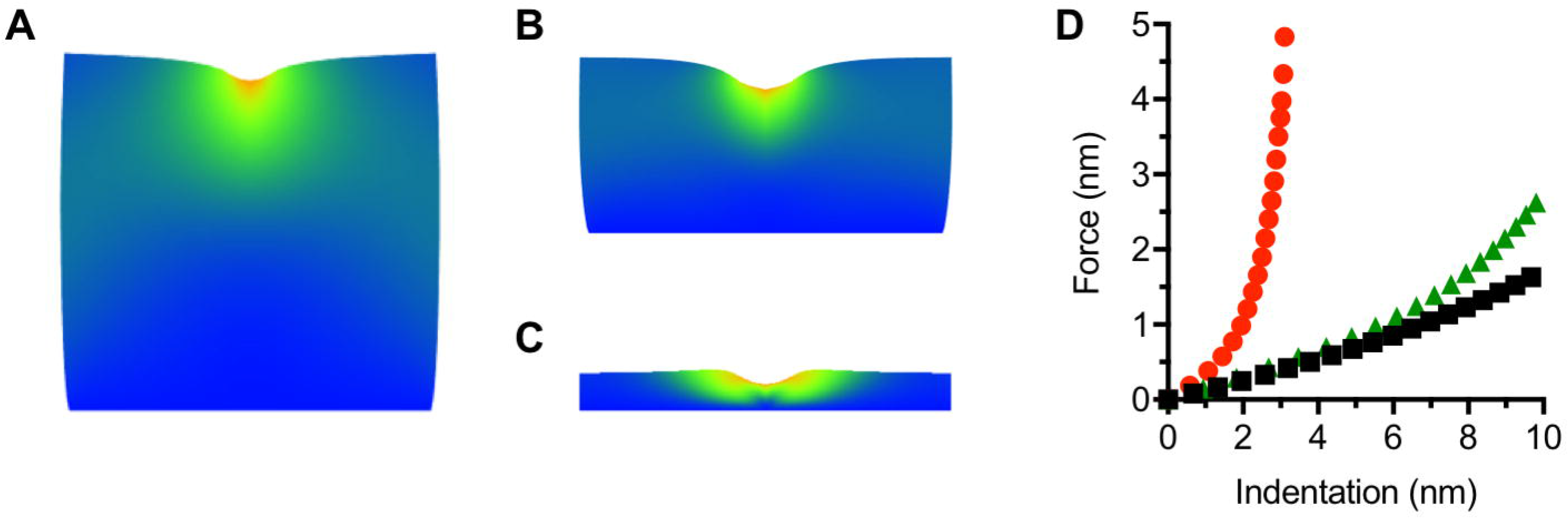
Finite element simulation of the deformation of a thin holdfast layer with Young's modulus given by 0.2 GPa for *h* = 120 nm (A), 60 nm (B) and 12 nm (C) thicknesses. The AFM tip is a cylindrical indenter with radius 13 nm. (D) The resulting force-indentation curves depend holdfast height, showing convergence for thicker samples at small indentation (black: 120 nm; green: 60 nm; red: 12 nm thickness respectively).

#### Surface molecular brush layer

The presence of the first regime in the F *vs*. D curves reveals a layer surrounding the stiff holdfast core, defined by a slow ramp in the force as a function of tip-sample distance during the approach (Fig. 2B and Fig. S1). This regime cannot be governed by electrostatic or ion-ion interactions, which have a shorter effective range (< 60 nm). The slow ramp is absent in the curve obtained on a clean mica substrate, indicating that the phenomenon is indeed specific to tip-holdfast interactions. We first hypothesize that the surface layer could be treated similarly to the holdfast core, with both regions described as elastic layers according to the Hertz model, albeit with different elastic moduli. However, creep compliance experiments performed at small dynamic loads and corresponding almost exclusively to indentations of the surface layer reveal significant creep at constant load (Fig. 4A). Thus, unlike the stiff core of solid-elastic nature, the soft surface layer behaves as a viscoelastic material. While the validity of Hertz’s linear elastic theory does extend to viscoelastic materials when contact area is only weakly dependent on the loading rate ^43^, we have no evidence that this condition holds, as the tip radius is generally smaller than the soft layer thickness. Moreover, attempts at using the Hertz model to describe both the surface layer and the bulk of holdfast as elastic materials with different elastic moduli do not yield satisfactory fits to the AFM data over its full range (Supplementary information).

**Fig. 4:**
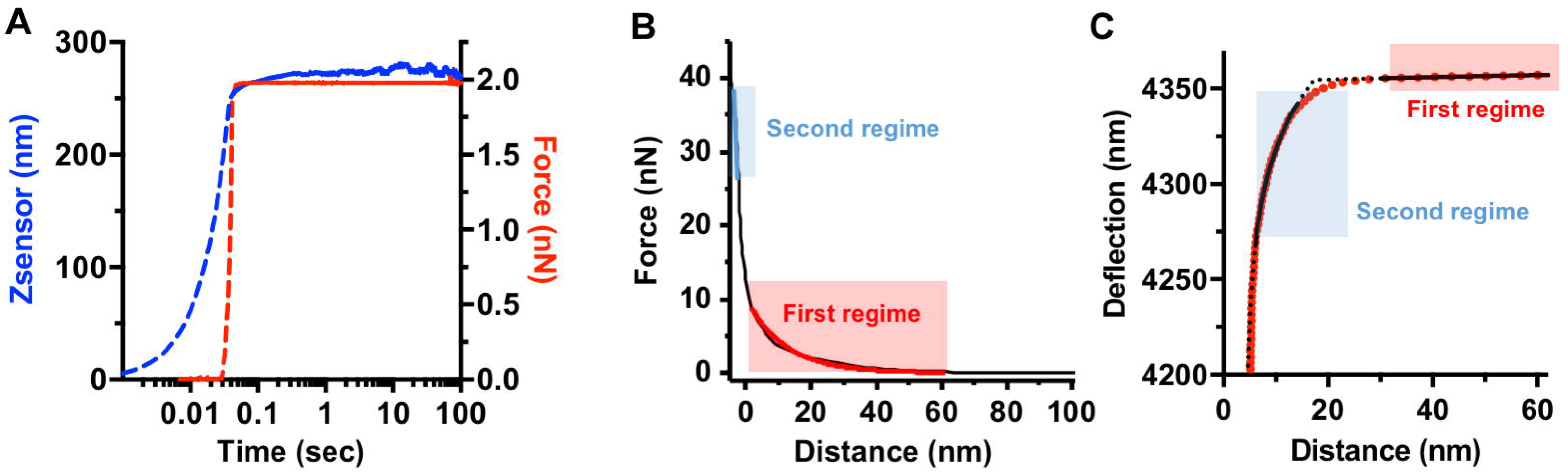
(A) Representative creep experiment data performed on a holdfast brush layer (incubated for 16 h) with a constant force of 2 nN and a dwell time of 120 sec. First, the tip indents the holdfast particle until the trigger force is reached (dotted lines). Once the probe reaches the set trigger force, the particle is deformed at a constant force along for 120 seconds (solid lines). The particle deformation over time is recorded as changes in the piezo extension Z (Zsensor channel, blue curve) and translated in Force (red curve). (B) Separate fits to brush layer model in the first regime (red line) and an effective, linear spring model in the second regime (blue line). (C) Simultaneous fits to brush layer and simple Hertz models in the first and second regimes, with fit parameters given by: AFM tip deflection far from the sample, d_0_ = 4.2 × 10^−9^ m; holdfast height, h = 42.5 × 10^−9^ m; holdfast surface, Z_0_ = 4.357 × 10^−6^ m; Young's modulus of the holdfast core, Ebulk = 0.348 × 10^9^ Pa; brush layer thickness, L_0_ = 279 × 10^−9^ m; brush layer density, T = 0.31 × 10^−17^ m^−2^. The fits in (E) and (F) exclude a transition region at the surface of holdfast where neither a linear elastic nor a brush layer model strictly holds.

Repulsive interactions similar in magnitude and range to those measured for the holdfast outer layer have been previously observed where biopolymers are present on the surface of different bacteria ^44,45^. As with holdfast, the magnitude and range of AFM interactions with the surface layer were much larger than those predicted by the Derjaguin, Landau, Verwey, and Overbeek theory ^44,46,47^. The repulsive entropic force between two surfaces coated with polymer chains was derived by Alexander ^48^ and de Gennes ^49^ and adapted by Butt et al. ^50^ to describe the force experienced by a bare AFM tip as it probes a polymer brush. The forces measured on the outer holdfast layer (first regime) are indeed comparable with those predicted by this brush layer model (with magnitude of force given by 2 - 10 nN, and range of force given by 50 - 100 nm, depending on the sample) (Fig. 2B). Hence, we explore the brush layer model as a quantitative description of the experimental data for the outer layer. We favored this approach over a continuous model of a cross-linked polymer mesh exhibiting a nonlinear stiffness profile ^51,52^. Such a continuous model would imply a well-defined holdfast/liquid interface extending around 50 - 100 nm from the substrate, while our data (Fig. S3) and others ^23,27,28^ show that the measured holdfast height is 40 nm in average, which corresponds to the stiff core regime in indentation experiments. In addition, our model of two regimes describing a stiff core and the polymer brush layer does not necessitate a discontinuity in the force-indentation curves, as a compression of the brush layer would result in transmission of some fraction of the applied force to the core layer, yielding a continuous response ^44-46,53,54^. Fig. 4B shows a typical data set, where the first and second regimes are separately fit to brush layer and linear spring models, respectively. In Fig. 4C, we also show simultaneous fits to the data in these regimes using the brush layer and simple Hertz models (Supplementary information). These fits exclude a transition region at the surface of holdfast, approximately 10-20% of the brush layer thickness, where the force response is neither purely elastic nor entropic in nature (Supplementary information), and consequently the Hertz and brush layer models do not strictly apply. Using both fitting methods, similar estimates are obtained for the equilibrium length (*L*_0_) and density (*Γ*) of the brush layer (Supplementary information, Table 1 and Fig. S4A-B and S4D-E), and the effective spring constant and Young's modulus are consistent (Supplementary information). Our results suggest that the holdfast consists of a stiff core with a Young’s modulus of 0.37 ± 0.18 GPa, decorated with a sparse biopolymer brush of 89.8 ± 5.2 nm thickness and ~10^17^ strands/m^2^ surface density (Table 1 and red box and whisker plots in Fig. S4). If we consider the holdfast core as a 40 nm radius hemisphere (Fig. S3), our results imply approximately 1,000 exopolymeric strands on the surface of a holdfast.

### Modifications of holdfast mechanochemical properties

#### Holdfast maturation

Previous studies suggested that holdfast cures over time, strengthening adhesion with the substrate ^21,27,28^. To further assess the putative effect of aging on the holdfast structure, we calculate core stiffness, equilibrium length (*L*_0_) and density (*Γ*) of the biopolymer brush layer after 16 h and 64 h incubation periods, from independent fits to the data in each of the brush layer and core regions, excluding the crossover regime (using the brush layer model fit for the first regime and extracting the slope of the linear dependence of the second regime, as described above). The biopolymer equilibrium length and density at 64 h show broader distributions with statistically different median values than those for 16 h incubation (Table 1, Fig. S2B and black box and whisker plots in Fig. S4). The results are consistent with a longterm re-arrangement of the brush layer, leading to a more compact biopolymer layer surrounding the core ^45^. Although the overall core stiffness does not statistically change between 16 and 64 h (Fig. S4), distributions are narrower at 64 h than at 16 h (Fig. S2 and S4), suggesting that the holdfast core becomes more homogeneous over time. Future experiments such as spatially resolved force curves will allow a better characterization the heterogeneous character of the holdfast, and its evolution over time.

#### Influence of ionic strength

As C. crescentus is an oligotrophic bacterium usually found in habitats with low concentrations of solutes ^17-19^, suggesting that holdfast mechanochemistry is optimized for such solutions. Modifying the ionic strength of a solution can greatly impact the conformation of bacterial polysaccharides ^55^. We previously showed that increasing the ionic strength has a strong negative impact on bulk adhesion of holdfast to surfaces ^28^ and here we investigate the influence of ionic strength on holdfast architecture (Fig. S5 and Table 1). Adding 10 mM NaCl modified the structure of the brush layer: its apparent length decreases by half, while the density of the polymeric strands doubles, suggesting a compaction of the overall brush layer ^45^. In addition, the holdfast core becomes twice as stiff in the presence of NaCl. These results are consistent with the 50% decrease in bulk adhesion of holdfast to glass in the presence of 10 mM NaCl ^28^, and argue that the holdfast adhesive is optimized for attachment in very low ionic strength environments such as those from which *Caulobacter* are typically isolated ^19^.

### Role of the Different Holdfast Components in Structure and Adhesion

#### Core and brush layer properties

In order to determine chemical factors responsible for the observed mechanical properties, we perform force spectroscopy measurements on holdfasts treated with three different enzymes: proteinase K (broad range peptidase) to digest proteins and/or peptides, DNase I to digest both single and doubled stranded DNA molecules, and lysozyme to digest 1,4 ß-linked NAG residues.

First, we study enzymatic effects on the core stiffness (Fig. 5A). The most marked differences with respect to untreated holdfast are observed after proteinase K treatment. The median stiffness value after proteinase K treatment is approximately three times larger than that of untreated holdfasts (Table 1). The width of the stiffness distribution, however, is significantly narrower than that of the control and all other treatments (Fig. 5A). This result suggests that peptide residues present in the holdfast core are crucial for heterogeneity and elasticity. Holdfast stiffness also drops by a third when treated with DNase I and lysozyme as compared to non-treated holdfasts, though the response is not as drastic as after proteinase K treatment, suggesting that both DNA and NAG molecules play a role in the constitutive properties of the holdfast core.

**Fig. 5:**
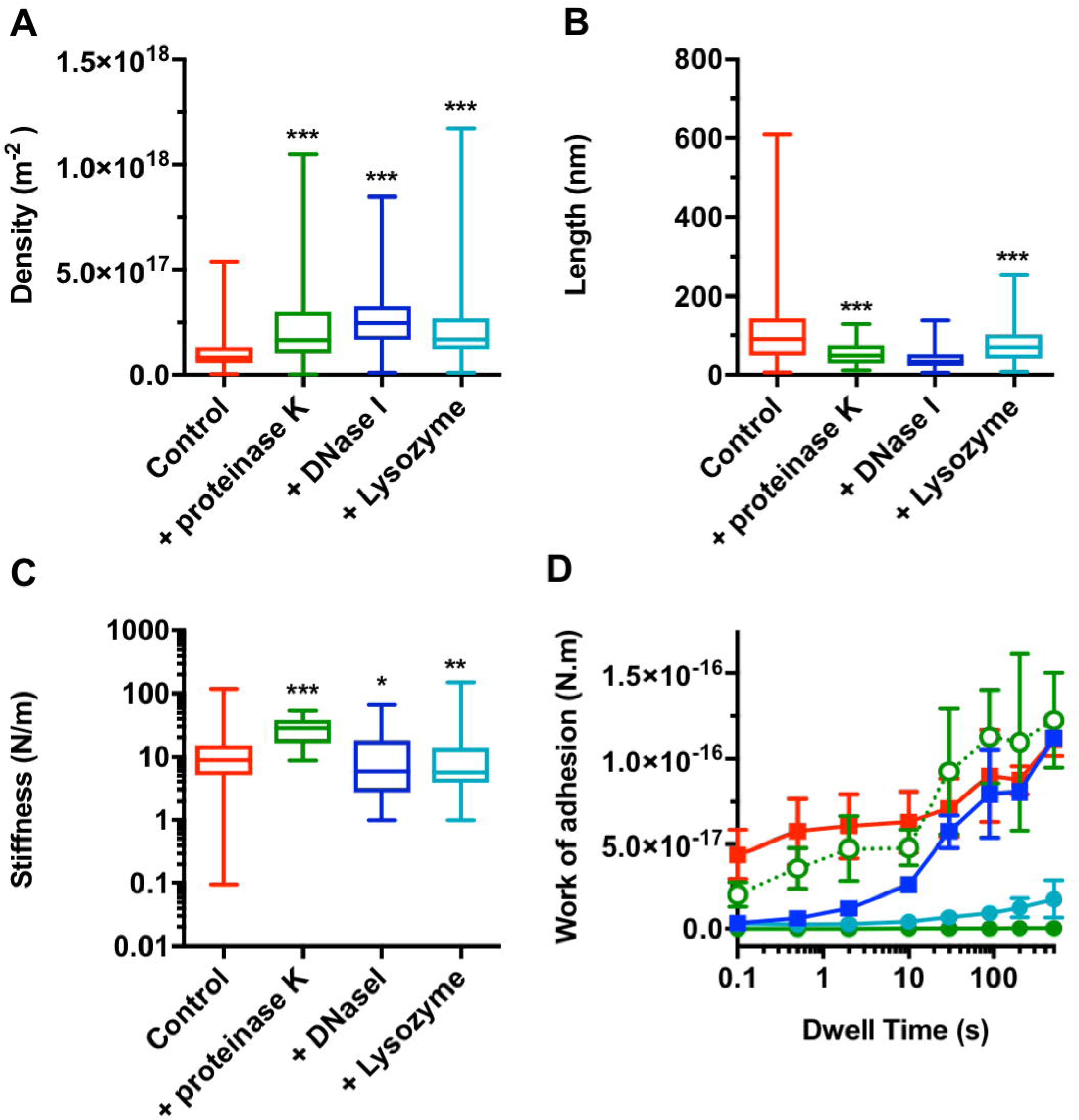
Post-enzyme treatment properties. Box and whisker plots of (A) core stiffness, (B) polymer brush equilibrium length, and (C) density of biopolymer brush layer for non-treated holdfast (red) and after treatment with proteinase K (green), DNase I (blue) and lysozyme (cyan). * p values < 0.025, ** p values < 0.01 and *** p values < 0.001 (Mann-Whitney unpaired t-tests). (D) Work of adhesion of non-treated (red), proteinase K (green), heat-inactivated proteinase K (green, open circles, dashed line), DNase I (blue) and lysozyme (cyan) treated holdfasts, as a function of dwell time (fixed trigger point = 500 pN). The error bars represent SEM.

None of the enzymatic treatments entirely remove the long range steric interactions between tip and holdfasts, but we observe quantitative changes in parameter values *L*_0_ and *Γ* between treated and non-treated holdfasts (Fig. 5B and 5C). The most marked changes in brush parameters occur after DNase I treatment. Equilibrium length *L*_0_ drops to nearly a third of that of the control (Table 1). At the same time, the apparent density increases, suggesting a form of condensation ^45^. A similar trend is observed for proteinase K treatment, though to a lesser extent (45% decrease in length). Taken together, our results suggest that DNA and peptide residues are important components of the biopolymer brush layer. The impact of lysozyme treatment on the brush layer is less pronounced, with only a 30% decrease in biopolymer length, suggesting that the NAG residues are not as important as DNA or peptide entities to the structure of the brush layer.

#### Dynamic Adhesion Force Spectroscopy

The nanoindentation experiments performed on enzyme-treated holdfasts reveal the presence of peptide and DNA entities in the holdfast. To determine whether these constituents play a role in the strength of adhesion, we perform dynamic adhesion force spectroscopy experiments on a holdfast-coated tip and a clean surface as previously described ^28^. Holdfast-coated AFM tips are first treated with different enzymes, and the strength of adhesion between the treated tips and clean mica is measured at different dwell times. Proteinase K treatment has a drastic effect on adhesion: a three order of magnitude decrease compared to the control sample was observed (Fig. 5D). This behavior was dependent on proteinase K enzymatic activity, since holdfast-coated tips incubated with heat-inactivated proteinase K exhibit the same adhesion as the non-treated control (Fig. 5D). This result suggests that peptide residues are crucial for initial holdfast adhesion strength. Exposure to lysozyme also leads to a large decrease in adhesion (roughly one order of magnitude), suggesting that NAG residues also participate in the adhesion of holdfast to surfaces (Fig. 5D). The results for DNase I treated samples are more intriguing (Fig. 5D): while adhesion strength is diminished for short dwell times (rapid contact between the holdfast and the mica), DNase I treated and non-treated tips behave similarly at dwell times higher than 10 sec. This suggests that DNA molecules are involved in the initiation of adhesion, but their contribution to adhesion is comparably less over time. Our results show for the first time that NAG residues are not the only components of the holdfast and highlight the presence of peptides and DNA molecules in the holdfast. These peptides and DNA molecules are not only part of the holdfast composition, they are also crucial for its adhesiveness.

#### Holdfast layer visualization

High-resolution AFM images of holdfast in air confirm the presence of long and thin structures associated with the holdfasts (Fig. S6A). However, due to the sparse (and possibly dynamic) nature of the brush, it is challenging to image it directly.

We use fluorescence microscopy to visualize the different components in the holdfast on whole cells and purified holdfasts (Fig. S6B). TexasRed Succinimidyl Ester (TRSE), which is an amine-reactive dye labeled holdfasts attached to cells (Fig. S6B), but staining of purified holdfasts with TRSE was not successful (Fig. S6B), suggesting this dye labels the holdfast anchor proteins present in the holdfast of wild-type cells, but absent in shed holdfasts ^56^. The same result is obtained when using fluorescent cysteine-reactive maleimide dye and we have yet to identify a method to label the peptide components in shed holdfasts revealed by protease treatment and AFM experiments. However, we successfully stain holdfast using the DNA dye YOYO-1, both on whole cells and purified holdfasts, confirming for the first time that DNA is a component of the holdfast (Fig. S6B). When we use wheat germ agglutinin lectin (WGA), to label the NAG residues ^25^ at the same time than YOYO-1, we can see colocalization of both labels (Fig. S6C). The fluorescence intensity profile plots strongly suggest that the DNA molecules are in the outermost layer of the holdfast, while the NAG residues are present in the inner layer (core). It is interesting to note that, when cells are clustered in rosettes (aka interact via their holdfast), the DNA layer seems larger than on isolated cells. When cells were treated with DNase I prior YOYO-1 labelling and imaging, only a faint labelling could be detected (Fig. S6C): the majority of the DNA molecules are easily accessible to the DNase I, suggesting that they are exposed, hence in the brush layer.

By imaging labelled cells using super resolution, structured illumination microscopy (SIM) (Fig. 6), we can clearly determine that NAG and DNA staining are spatially segregated. In addition, it seems that holdfast-holdfast interaction is mediated by the DNA molecules. Indeed, in rosettes, we can detect several NAG patches, probably corresponding to the cores of several holdfasts, interconnected by DNA molecules. The origin of the DNA present in the holdfast is still elusive at this stage, namely as an intrinsic component of the secreted holdfast, or as extracellular DNA (eDNA) released in the culture when bacteria lyse. We previously showed that eDNA released during cell death can interact with holdfast and prevent adhesion ^57^, suggesting the intriguing possibility that the brush layer DNA is an intrinsic part of the holdfast that can be bound by eDNA released by cell death to inhibit adhesiveness.

**Fig. 6:**
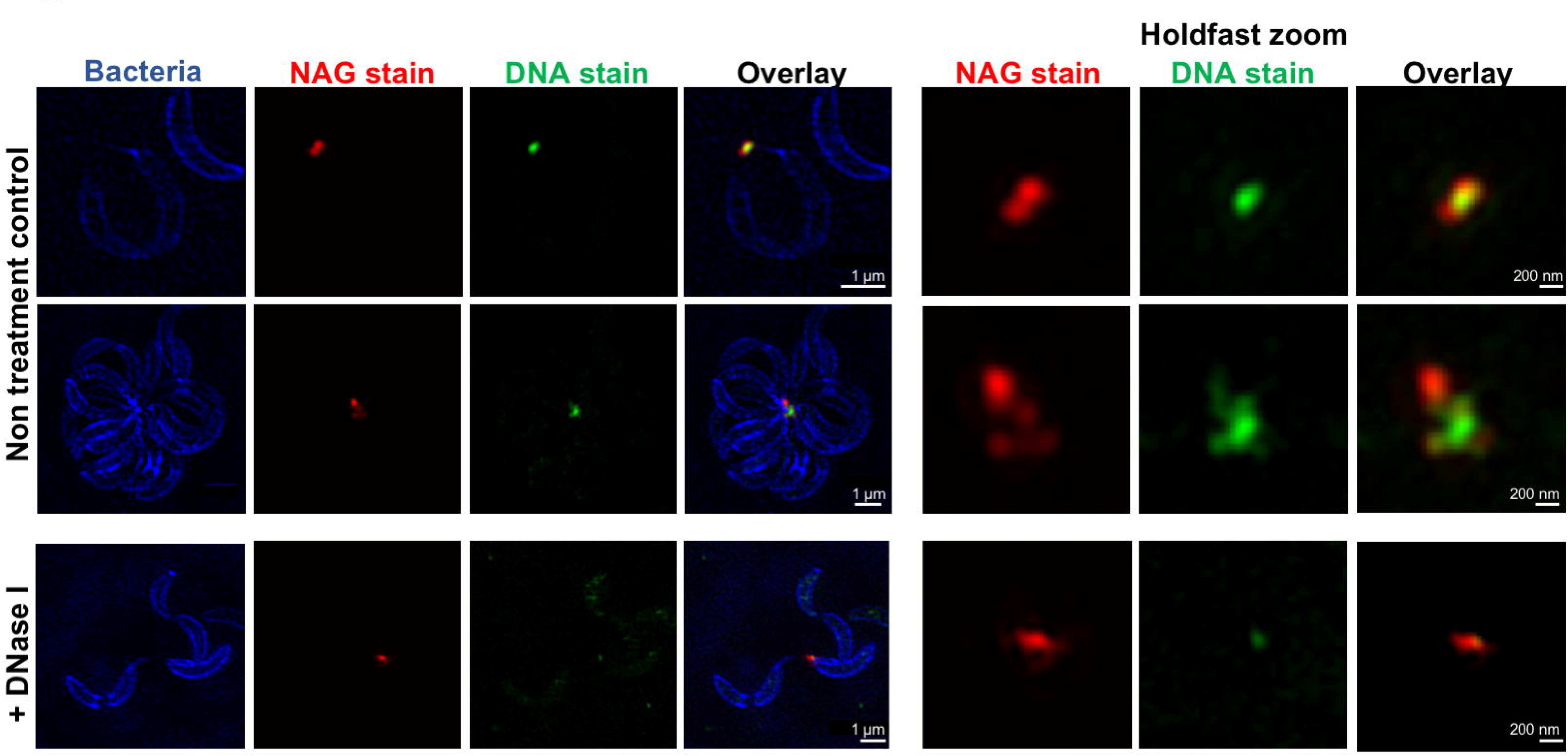
Structured Illumination Microscopy images of *C. crescentus* cells attached by their holdfast. Cells are stained using HADA (to label peptidoglycan), while NAG and DNA residues in the holdfast are labeled using AF-594 WGA (red) and YOYO-1 (green) respectively. Representative Z-stack images are presented. For the holdfast zoom panels, the average intensity for the entire stack for the green and red channels are projected together and merged in a single image.

## Conclusion

The holdfast has impressive versatility in strongly binding wet surfaces of variable roughness and hydrophobicity ^28^. While our previous understanding of holdfast as a simple NAG polymer was hard to reconcile with its adhesive strength and versatility, detailed determination of its composition by traditional methods remained elusive due to its insolubility and synthesis in small amounts. Here, using AFM dynamic force approaches, new structural and local chemical characteristics of the holdfast are unveiled providing significant insight into its properties. Our data support a model where the holdfast is an organized two-layer system. The remarkably stiff core, whose mechanical characterization suggests that it becomes more homogeneous over time, is surrounded by a relatively thick and sparse polymer brush layer. The properties of the newly identified brush layer are strongly affected by proteinase K and DNase I treatment, indicating that DNA and peptides play an important role in its structure. While peptides appear to be crucial for holdfast adhesion strength at all stages, DNA appears to be involved only in the initial adhesion step.

Previous work showed that the kinetics of adhesive bond formation vary with surface chemistry and roughness, suggesting that initial adhesion is substrate-dependent ^28^. However, adhesion forces on different surfaces converge to similar values over time, suggesting that once the initial substrate-dependent intramolecular rearrangement is complete, adhesion is strengthened in a substrate-independent manner. The multi-regime structure of the holdfast reported here suggests a hypothetical model for holdfast adhesion wherein flexible brush layer fibers readily explore a surface, forming the first, initially weak contacts. Compaction of the brush layer would serve to bring the holdfast core in contact with the surface. Stiffening of the core from its secreted fluid state to a hardness comparable to epoxy glues allows for an overall curing, whereby an applied force is distributed among the surface bonds and the load-dependent rate of detachment is reduced. A clear advantage of this complex structure and mechanochemistry is that each component can be modified to yield a diversity of adhesive strategies adapted to various environments. Other bacteria closely related to C. crescentus generate similar nanoscopic adhesive structures ^4,15,16,19,25^. Their habitats range from saltwater to freshwater to the rhizosphere, necessitating different conditions for effective adhesion. This framework can be used to characterize these bacterial bioadhesives, as well as a benchmark for the development of a versatile, new family of synthetic adhesives.

## Methods

### *Caulobacter crescentus* purified holdfast sample preparation

*C. crescentus* CB15 Δ*hfaB* (YB4 2 51) ^56^ was grown in Peptone-Yeast Extract medium ^19^ at 30°C. In this strain, the gene encoding HfaB, a key protein for holdfast anchoring to the bacterial cell, is deleted, yielding a shedding phenotype: the holdfast is produced and exported outside the cell, but fails to stay attached to the cell body ^56^. 100 μl of a *C. crescentus ΔhfaB* culture (A 600 nm = 0.3 − 0.5) was spotted on a freshly cleaved piece of mica and incubated at 30°C in a humid chamber. After 16 h incubation in PYE medium, the mica surface containing shed holdfasts was carefully rinsed with sterile dH_2_O to achieve a low ionic strength condition typical of the oligotrophic environments from which *C. crescentus* is typically isolated. Indeed, the ionic strength of river and lakes where *Caulobacter* can be readily found attached to surfaces is very low and typically ranges between 1 and 5 mM^17–19^. In addition, *Caulobacter* is readily found in potable and filtered water systems. These conditions are consistent with conditions of previous studies ^28^. This step removes the bacterial cells, while the shed holdfasts stay attached to the surface ^28^. 100 μl of sterile dH2O was placed on top of the holdfasts attached to the mica prior to AFM analysis. For 64 h incubation time samples, bacteria were removed from the surface after 16 h, as described above, and holdfasts attached to the mica were incubated in water in a humid chamber at room temperature for an additional 48 h. For holdfast enzymatic treatments, holdfasts were treated with the following enzymes: proteinase K, a non-specific serine protease (10 μg/ml); lysozyme, hydrolase specific for 1,4 ß-links of NAG polymers (20 μg/ml); and DNase I, endonuclease cleaving single and double-stranded DNA (10 μg/ml). Enzymatic treatments were performed for 1 h at room temperature and were done on holdfasts bound to mica surfaces or immobilized to AFM tips.

### Atomic Force Microscopy experiments

AFM measurements were performed using a commercial Cypher AFM instrument (Asylum Research) operating at room temperature.

#### Holdfast Indentation and Creep Compliance Experiments

Samples were imaged using tapping mode AFM. The AFM probe was driven near the first resonance frequency of its flexural mode and then engaged on the sample. The cantilever spring constant and quality factor of the first flexural mode was calibrated using the thermal noise method in liquid ^58^. Excitation frequency was chosen from the peak of the tuning curve, where the phase lag became 90°.

To corroborate the elasticity behavior of the holdfast core over time, we performed creep compliance experiments. In these experiments, the holdfast is deformed and the change of deformation over time, under constant load is measured ^33^. First, a holdfast was centered in the scan area after imaging. Then, the AFM tip was moved on the top of the particle and a *F *vs*. Z* curve was acquired with a trigger force around 40 nN to probe the holdfast core or 2 nN to probe the brush layer and a loading rate from 6.25 μm/sec (Fig. 2A). During this first stage, the tip indented the particle until the selected trigger force was reached. Once the tip reached the set trigger force, the particle was deformed at a constant force using a dwell time of 180 sec. The *Z*-sensor channel was recorded to monitor the response of continuous deformation upon constant loading force. After each measurement, a new image was recorded to compare the holdfast morphology before and after indentation. Images were processed using the WSxM software ^59^.

For nanoindentation experiments in liquid environment, a high resolution image (360 nm × 360 nm, 128 points) of a given holdfast was recorded in Amplitude Modulation AFM (AM-AFM), in order to check the morphology and locate the center of the holdfast. We used a soft silicon nitride microcantilever (Olympus RC800) with a tip radius of 15 nm, a nominal spring constant of 0.3 − 0.4 N/m and a resonance frequency of 69 kHz (~ 24 kHz in liquid) that was excited at an amplitude A_free_ ~ 4 nm (A/A_ratio_ = 0.85 − 0.90). Once the holdfast was centered in the scan area, the AFM tip was moved on top of the particle and a set of 3 − 8 force-displacement (*F *vs*. Z*) curves (40 nN trigger force) was recorded at different locations within the same holdfast. For trigger forces < 5 nN, the holdfast exhibited a linear elastic behavior. Force *vs*. indentation curves (F *vs*. Ind) were calculated according to the spring constants of the cantilever and to the measurements of cantilever deflection on mica and on holdfast ^60^ (Fig. S1).

#### Dynamic Adhesion Force Spectroscopy Experiments

Dynamic Adhesion Force data were obtained from the interaction of a clean surface (freshly cleaved mica) and holdfast immobilized on the AFM tip in a liquid environment, as described previously ^28^, using silicon nitride gold covered microcantilevers (MicroMasch HQ: CSC38), with a tip radius of □ 30 nm, nominal spring constant of ~ 0.09 N/m.

First, we characterized the tip before each experimental set data: a F *vs*. Z curve was performed on a clean mica surface with a clean AFM tip. The retraction curve showed a characteristic adhesion between tip and mica. Then, holdfast was immobilized on the same tip. To do so, a holdfast bound on the mica surface was first localized by imaging; then some holdfast material was transferred to the AFM tip by several contacts between the holdfast on the surface and the tip, using a dwell time of 90 sec a trigger force of 500 pN. Our choice of a trigger force is based our previous AFM study of holdfast ^28^ in which we tested adhesion forces of purified holdfasts using different trigger points spanning one order of magnitude on either side of this value (from 50 to 5000 pN) and showed that for trigger forces between 250 pN and 5 nN, the work of adhesion and maximal force measurements remained constant. To confirm the transfer, a second F *vs*. Z curve was performed on clean mica. Only holdfast covered tips which presented stronger adhesions compared to clean tips were further considered for our analysis. Once holdfast material was immobilized on the AFM tip, a new freshly cleaved mica surface was used for F *vs*. Z curves, using different dwell times (from 0.1 to 500 sec) and a trigger force of 500 pN. Data were analyzed as previously described ^28^.

#### Nanoindentation Data Analysis

For all AFM indentation experiments, 3 − 8 different measurements were taken at different locations in a single holdfast. For 16 h incubation samples, 86 holdfasts were measured in 10 independent experiments, yielding to a collection of 310 F *vs*. Z curves. For 64 h incubation samples, 59 holdfasts were measured in 8 independent replicates, yielding to a collection of 298 F *vs*. Z curves. For enzyme-treated holdfasts, 17, 34 and 25 different holdfasts were measured in at least 2 independent replicates for proteinase K, DNase I and lysozyme treatments respectively. To characterize the interactions between the AFM tip and the holdfast, we converted the *F vs. Z* curves into force *vs*. distance tip-sample curves (F *vs*. D; gap distance, *D = Z − Z_0_ − d*), where *Z* is the piezo displacement, *Z_0_* is the point to contact and d is the average cantilever normal deflection, as routinely performed in AFM spectroscopy for rigid surfaces ^60,61^ (Fig. S1), using a home script based on the Igor Pro software.

We observed approximate linear behavior in the limit of large forces and characterized this approximate linear response of the holdfast bulk with an effective spring constant, *k*. We defined the contact point *Z_0_* between the tip and sample as the distance where the *F vs. D* curve departs from the linear behavior ^44,50,60,61^. Holdfast deformation is negligible under forces smaller than 5 nN (Fig. S1 E). The strength and range of the tip-holdfast interaction forces measured in our experiments (forces close to 5 nN and a tip-sample interaction gap measured range > 50 nm) suggest that forces other than the classical van der Waals and electrostatic forces are involved, as described previously in other biological systems ^44,45^. Indeed, these previous AFM measurements have described the contribution of steric forces in various bacterial systems, where a biopolymer brush layer covering the bacterial cell surface is responsible for long-range interaction forces ^44,45,62^.

The AFM data over the slow ramp range (2 − 60 nm) were fitted using the model developed to describe the interaction between an AFM tip and a grafted polymer surface (*F_brush_*) ^50^, based on the theoretical work by Alexander ^48^ and de Gennes ^49^:

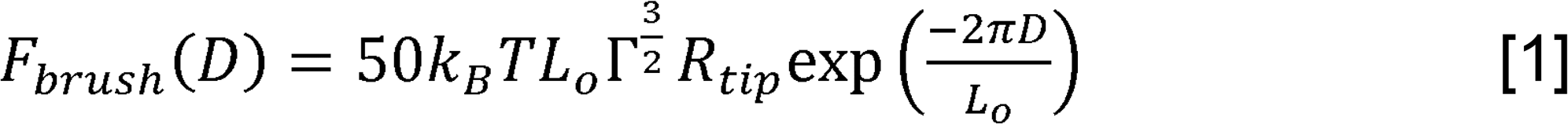

where *T* is temperature, *k_B_*, is the Boltzmann constant, *L_0_* is the equilibrium length of polymer brush, *R_tip_* is the tip radius (15 nm), *Γ* is the grafted polymer density and *D* is the distance between tip and sample.

Using equation [1] and Igor Pro software, we estimated the equilibrium length *L_0_* and the density *Γ* of the holdfast brush layer. Only curves with an acceptable fit (error < 10%) were considered for further analyses.

### Finite Element Calculations

The finite element method was employed to simulate the indentation on films of different thicknesses on a stiff substrate. The three substrates were chosen to be 120 nm at the base. We imposed the homogeneous Dirchlet boundary condition at the bottom of the film. We applied a force at the top of the film and slowly increment the force at each iteration. We considered the indenter as a cylindrical end with a radius of 6.5 nm. The contact between the indenter and the film was assumed to be frictionless and the body force was assumed to be zero. For all three cases, the Young's modulus was set to be 0.2 GPa and the Poisson ratio was set to be 0.45. Moreover, a linear constitutive law was assumed.

### Simultaneous fits to Hertz and brush layer models

For a spherical tip and for small indentation, the Hertz model predicts the following relationship between force and indentation:

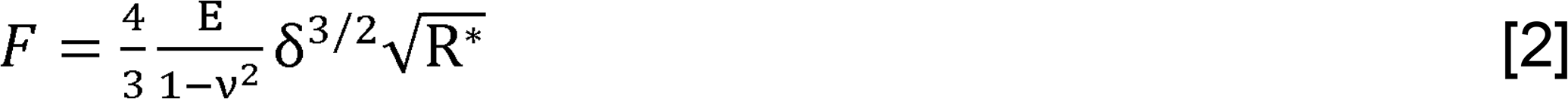

where *E* is the Young’s modulus of the holdfast; *v* is the Poisson ratio, taken to be 0.5 here, assuming incompressibility of the holdfast, and *R* = R_tip_ R_sample_/ (R_tip_ + R_sample_*) where for typical sample radii, *R* ≈ R_tip_*. The sample indentation, δ, is given:

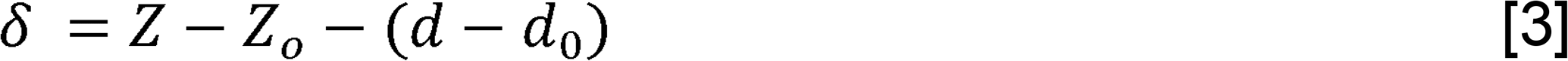

where *d* is the cantilever deflection, *Z* is the piezo height, *d_0_* is the deflection of the cantilever far from the sample, and *Z_0_* represents the piezo displacement for which the cantilever touches the surface of the holdfast core. The force was obtained from the cantilever deflection using the cantilever spring constant, *k_c_: F = k_c_ (d − d_0_*).

We extracted parameters describing holdfast material properties from the raw AFM data by performing simultaneous least-squares fits to Hertz and brush layer models (Fig. 2 and Table 1). Specifically, with the cantilever deflection given by d, and piezo height, Z, we plotted Z − d vs. d, and fitted the data in selected regions to functional forms describing the bulk of the holdfast and a surface brush layer, as described below. We chose to plot the data this way, as the fit procedure is most robust for the portion of the data at large d, where Z - d varies slowly with d, allowing reliable extraction of the modulus of elasticity, E.

We followed two approaches to fit the AFM data simultaneously to the Hertz and brush layer models: in the first approach, we fitted the brush layer model all the way to the surface (supplementary information), where we allowed the boundary between these regions to float as a fit parameter in the least-squares minimization process. For values of d greater than this value, the fit function is the Hertzian form, given by

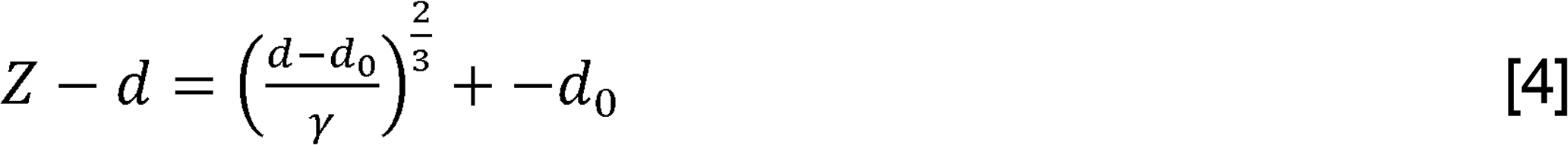

where the Young’s modulus is obtained from the fit parameter γ as

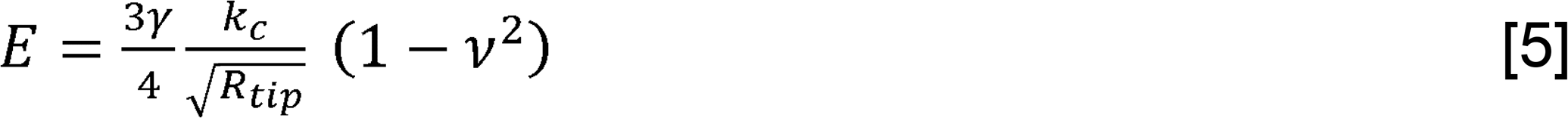

For values of *d* less than the boundary, the fit function represents a description of the brush layer

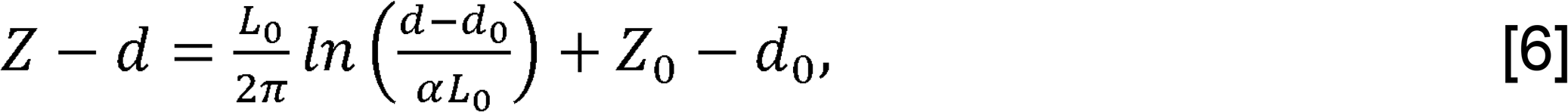

where brush layer density is obtained from the fit parameter α as

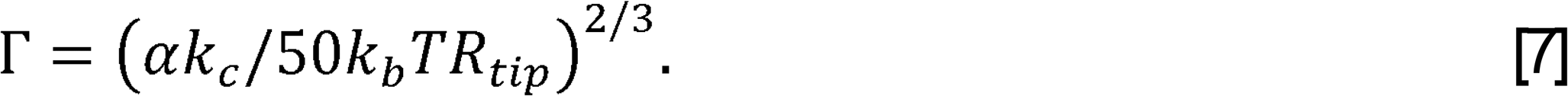

In the second approach, we excluded the transition region, corresponding to *D/L_0_* less than ~ 0.1 −0.2, from the brush layer fitting region, wherein the force is not strictly entropic. In the SI, we show that the two approaches yield consistent results for the values of fitted parameters to within an order of magnitude (Table 1 and Fig. 4). However, we emphasize that in the transition region between the bulk and surface layer, the brush layer is highly compressed and neither model constitutes an accurate physical description.

### Fluorescence Microscopy

Holdfast were fluorescently labelled and visualized on whole *C. crescentus* CB15 Wild-type cells. Exponential cultures (OD_600_ = 0.4 – 0.7) grown in PYE medium ^19^ were stained using 0.5 μg/ml AlexaFluor 647 conjugated Wheat Germ Agglutimin lectin (AF647-WGA, Molecular Probes) and 1 μM YOYO-1 DNA stain (Molecular Probes) and incubated 5 min incubation at room temperature. WGA specifically binds to the NAG residues present in the holdfast ^25^, while YOYO-1 is a cell-impermeant dye that has a high affinity for dsDNA molecules. For TexasRed Succinimidyl Ester (TRSE, amine reactive dye, Molecular Probes) staining, cells were mixed with 5 μg/ml dye (1/1000 dilution in 100 mM NaCO_3_ buffer, pH8) and incubated for 20 minutes at room temperature, before being washed 3 times by centrifugation (3,000 g for 2 min) and resuspended in dH_2_O. 1 μl of labelled cells was spotted onto a 24×60 microscope glass coverslip and covered by an agarose pad (1% in water). Samples were imaged by epifluorescence microscopy using an inverted Nikon Ti-E with a Plan Apo 60X objective, an Andor iXon3 DU885 EM CCD camera and Nikon NIS Elements imaging software.

### Transmission electron microscopy (TEM)

Exponentially grown *C. crescentus* CB15 Wild-type were spotted onto Formvar-coated, carbon film-stabilized copper grids (Electron Microscopy Sciences) and incubated for 1 h. Each grid was washed with dH_2_O, negatively stained with 7.5% uranyl magnesium acetate for 5 min, and washed five times with dH_2_O again. Imaging was performed with a Jeol J EM-1010 transmission electron microscope set to 80 kV.

### Structured Illumination Microscopy (SIM)

For SIM imaging, we used HADA, a blue fluorescent D-amino acid ^63^ to label *C. crescentus* CB15 peptidoglycan, YOYO-1 (Molecular Probes) to label dsDNA present in the holdfasts and AF594-WGA (Molecular Probes) to label NAG residues present in the holdfast. 1 ml of *C. crescentus* CB15 was grown to exponential phase in PYE and labelled with 1 μM YOYO-1 and 0.5 μg/ml AF594-WGA for 5 min at room temperature. Cells were washed 5 times by centrifugation (3,000 g for 2 min) in 1 ml PYE to remove all traces of unbound YOYO-1 and AF594-WGA. 1 mM HADA was added to the washed cells in 1 ml PYE. After 25 min of incubation at 30°C under constant shaking (200 rpm), cells were washed again 3 times by centrifugation (3,000 g for 2 min) to remove unbound HADA and resuspended in 50 μl water. 1 μl was spotted onto an 24x50 microscope glass coverslip and covered by an agarose pad (1% in water) before imaging. Because of the multiple wash steps during the labeling process, all cells harboring a holdfast are arranged in rosettes, clusters of cells interacting together via their holdfasts, and we cannot detect isolated cell with a holdfast.

*Z*-series images of cells harboring a holdfast images were acquired on a DeltaVision OMX 3D-SIM super-resolution system (Applied Precision Inc) equipped with an inverted 1.4 NA Olympus 100 × oil objective. Images were processed (deconvolution and alignment) using Softworx imaging software.

## Acknowledgments

We thank Yen Pang Hsu and David Kysela for their input on fluorescent microscopy, and the members of the Brun laboratory for critical comments on the manuscript. This work was partially supported by the U.S. Army Research Office under Award W911NF-13-1-0490 (to B.D. for contact mechanics modeling) and by National Institutes of Health grants GM077648 to Y.V.B. and B.D. and R35GM122556 to Y.V.B. SIM and TEM experiments were performed in the Indiana University Light Microscopy Imaging Center and Electron Microscopy facility at Indiana University.

## Author contributions

M.H.P, B.D., Y.V.B. and C.B. designed the research. M.H.P. and C.B. performed the experiments. S.S., Y.H., R.T. and B.D. performed the theoretical analysis. M.H.P., S.S., B.D., Y.V.B. and C.B. analyzed the data. M.H.P., S.S., B.D., Y.V.B. and C.B. wrote the manuscript.

